# The lysine demethylase dKDM2 is non-essential for viability, but regulates circadian rhythms in *Drosophila*

**DOI:** 10.1101/291070

**Authors:** Yani Zheng, Yongbo Xue, Xingjie Ren, Xiao-Jun Xie, Mengmeng Liu, Yu Jia, Xiao Li, Ye Niu, Jian-Quan Ni, Yong Zhang, Jun-Yuan Ji

## Abstract

Post-translational modification of histones, such as histone methylation controlled by specific methyltransferases and demethylases, play critical roles in modulating chromatin dynamics and transcription in eukaryotes. Misregulation of histone methylation can lead to aberrant gene expression, thereby contributing to abnormal development and diseases such as cancer. As such, the mammalian lysine-specific demethylase 2 (KDM2) homologs, KDM2A and KDM2B, are either oncogenic or tumor suppressive, depending on specific pathological contexts. However, the role of KDM2 proteins during development in the whole organisms remains poorly understood. Unlike vertebrates, *Drosophila* has only one KDM2 homolog (dKDM2), but its functions *in vivo* remain elusive due to the complexities of the existing mutant alleles. To address this problem, we have generated two *dKdm2* null alleles using the CRISPR/Cas9 technique. These *dKdm2* homozygous mutants are fully viable and fertile, with no developmental defects observed under laboratory conditions. However, the *dKdm2* null mutant adults display defects in circadian rhythms. Most of the *dKdm2* mutants become arrhythmic under constant darkness, while the circadian period of the rhythmic mutant flies is approximately one hour shorter than the control. Interestingly, opposite defects are observed when dKDM2 is overexpressed in circadian pacemaker neurons. Taken together, these results demonstrate that *dKdm2* is not essential for viability; instead, dKDM2 protein plays important roles in regulating circadian rhythms in *Drosophila*. Further analyses of the molecular mechanisms of how dKDM2 and its orthologs in vertebrates regulate circadian rhythms will advance our understanding of the epigenetic regulations of circadian clocks.

## Introduction

Covalent histone modifications, particularly methylation and acetylation at lysine (K) residues on the N-terminal tails of histones H3 and H4, play fundamental roles in epigenetic regulation of gene expression in eukaryotes. The addition or the removal of methyl groups to lysine residues is dynamically modulated by specific lysine methyltransferases (KMTs) and lysine demethylases (KDMs), respectively (Shi and Whetstine 2007; Black *et al.* 2012). These enzymes are highly conserved in the evolution of eukaryotes (Zhou and Ma 2008; Zhang and Ma 2012; Qian *et al.* 2015). Extensive efforts have been devoted to elucidating the molecular mechanisms of how these enzymes catalyze reactions towards specific lysine residues (Shi and Whetstine 2007; Black *et al.* 2012). Based on the chemical reactions that they catalyze, the KDMs are classified as two major families: the lysine-specific histone demethylase (LSD) family, including LSD1 (also known as KDM1) and LSD2, and the Jumonji C (JmjC) domain family demethylases, containing more than 20 KDMs in mammals (Shi and Whetstine 2007; Shi and Tsukada 2013). The LSD family demethylases can only demethylate mono- and dimethylated (me1 and me2) lysine residues through an FAD-dependent amine oxidase reaction; while the JmjC domain family demethylases can demethylate me1, me2, and trimethylated (me3) lysine residues through a 2-oxoglutarate-Fe(II)-dependent dioxygenase reaction (Shi and Whetstine 2007; Cloos *et al.* 2008; Shi and Tsukada 2013). Since the initial discovery of LSD1 (Shi *et al.* 2004), we have witnessed remarkable progress in the identification and characterization of these demethylases in diverse organisms, particularly in their substrate specificities and their effects on regulating transcription at single gene and genome-wide levels. However, the *in vivo* functions and regulations of these enzymes in developmental, physiological, and pathological contexts are still not fully understood (Nottke *et al.* 2009; Dimitrova *et al.* 2015).

Based on their sequence homology and their specificities in removing the methylation marker on lysine residues, the JmjC KDMs (KDM2-7) are further grouped into five clusters, including KDM2/7, KDM3, KDM4, KDM5, and KDM6 (Klose *et al.* 2006; Cloos *et al.* 2008). In vertebrates, the KDM2 subfamily of KDMs consists of KDM2A and KDM2B, and studies in the past decade have linked mutations of these KDM2 paralogs to a number of human cancers (Klose *et al.* 2006). For examples, KDM2A has been reported to play an oncogenic role in breast cancer (Liu *et al.* 2016); overexpression and the oncogenic role of KDM2B has also been observed in acute myeloid leukemia (He *et al.* 2011), pancreatic cancer (Tzatsos *et al.* 2013). However, KDM2A appears to play a tumor suppressive role in colon cancer (Lu *et al.* 2010), and similarly, a tumor suppressor role of KDM2B has been implicated in lymphoma and brain cancer (Suzuki *et al.* 2006; Frescas *et al.* 2007). Thus it seems that the role of KDM2A and KDM2B in tumorigenesis is dependent on different types of cancers. Elucidation of the *in vivo* functions of these enzymes is key to understand how misregulation of KDM2 proteins contributes to these different cancers.

To address this problem, we have performed developmental genetic analyses of *Kdm2* mutants in *Drosophila*, because of its relative simplicity compared to mammals. *Drosophila* also provides sophisticated genetic tools to dissect the function and regulation of genes of interest. *Drosophila* has 13 KDMs, with only one conserved KDM2 ortholog, dKdm2 (Lagarou *et al.* 2008; Kavi and Birchler 2009; Zheng *et al.* 2014; Holowatyj *et al.* 2015). Based on analyses of 13 mutant alleles of *dKdm2* gene, we have concluded that *dKdm2* is not required for normal development in *Drosophila* (Zheng *et al.* 2014). This notion is unexpected for the following reasons: first, dKdm2 has been identified as a subunit of the dRING-associated factor (dRAF) complex, a Polycomb group (PcG) complex that was proposed to play critical roles in coordinating trans-histone regulation by replacing the active H3K36me2 marker with a repressive mono-ubiquitinated H2A K118 (H2AK118ub or H2Aub) marker (Lagarou *et al.* 2008). Second, dKdm2 was shown to demethylate H3K36me2 but not H3K36me1/3 or H3K4me3 in *Drosophila* S2 cells (Lagarou *et al.* 2008), but depletion of dKDM2 in *Drosophila* larvae specifically increased the levels of H3K4me3 but not H3K36me2 (Kavi and Birchler 2009). Third, it has been reported that the dRAF complex is required for producing H2AK118ub in *Drosophila* (Lagarou *et al.* 2008), yet a recent report demonstrates that the bulk of H2AK118ub is produced by the RING1-L(3)73Ah complex instead (Kahn *et al.* 2016).

A major concern in our previous genetic analyses is that all existing mutant alleles of the *dKdm2* locus have their limitations: the transposon insertion lines are either hypomorphic or fail to affect the mRNA and protein levels of dKDM2, while the three deficiency lines uncover the *dKdm2* locus disrupt both *dKdm2* and its neighboring genes (Zheng *et al.* 2014). Having a molecularly defined *dKdm2* null allele that only disrupts the *dKdm2* locus is thus required to resolve the aforedescribed contradictory observations, and will enable us to draw stronger conclusions about the *dKdm2* null phenotypes and its molecular functions.

Using the CRISPR-Cas9 technique, we have generated two null alleles that only disrupt *dKdm2* but not its neighboring genes. Our analyses of these two null alleles demonstrate that *dKdm2* gene is indeed not required for normal development, viability, or fertility. The methylation states on H3K36 and H3K4 are mildly elevated in the homozygous mutant larvae, but no effect on H2Aub was observed in these mutants. In contrast, overexpression of wild-type dKDM2 in larvae or wing discs only reduced H3K36me2 and to a less extent H3K36me1, but not the levels of H3K36me3. Although the *dKdm2* homozygous mutants are fully viable, the mutant flies are defective in circadian rhythms: they lose circadian rhythms and have a shorter circadian period than the control. Consistently with this, overexpression of *dKdm2* in circadian neurons increases the circadian period. Taken together, our results show that dKDM2 regulates circadian rhythms, but does not play any essential roles during normal development.

## Materials and Methods

### Fly strains

*Drosophila* strains were maintained on standard cornmeal-molasses-yeast food at 25°C, and the *w*^*1118*^ line served as the control. The following Gal4 lines were obtained from the Bloomington *Drosophila* Stock Center: *engrailed-Gal4* (*en-Gal4*; BL-1973), *Sgs3-Gal4* (BL-6870), and *ubiquitin-Gal4* (*Ubi-Gal4*; BL-32551).

### Generation of the *UAS-dKdm2*^*+*^*-eGFP* transgenic line

The *dKdm2* cDNA was subcloned to the pENTR vector using the pENTR/D-TOPO Cloning kit (Invitrogen, K240020). The pTWG vector (from the DGRC #1076: https://dgrc.bio.indiana.edu/product/View?product=1076) was used as the destination vector and the *att*L x *att*R reaction was mediated by LR Clonase II enzyme mix (Invitrogen, 11791-020), resulting the ‘*pUASt-dKdm2*^*+*^*-eGFP*’ transgenic vector. After injecting the vector into *w*^*1118*^ embryos, the *UAS-dKdm2*^*+*^*-eGFP* transgenic lines were established by standard fly genetics.

### Generation and validation of the *dKdm2* null alleles *dKdm2*^*1*^and *dKdm2*^*2*^

Two *dKdm2* null alleles were generated using the *Drosophila* germline specific CRISPR/Cas9 system (Ren *et al.* 2013; Ren *et al.* 2014). Four sgRNAs were designed to target coding region of *dKdm2* (NO3: AGCTGGTAGAGCGAAGGCGG, NO4: GAGGATGGCGAGGGGACGCG, NO5: TCTTCAAAGTTCGCGCAGGC, NO6: TACATTGCTGCCGCCCCCGG) and cloned into the U6b-sgRNA plasmid. To generate *dKdm2* null alleles, sgRNA plasmids (sgRNA3, 4, 5 for *dKdm2*^*1*^, sgRNA4, 5, 6 for *dKdm2*^*2*^) were injected into *nos-Cas9 Drosophila* embryos. All adult flies developed from injected embryos were crossed with the “*y*^*1*^ *sc*^*1*^ *v*^*1*^; *+; Dr*^*1*^ *e*^*1*^*/TM3 Sb*^*1*^” stain, and null alleles were screened in F1 generation with single crossing and *Drosophila* wing genomic DNA PCR. *dKdm2*^*1*^ and *dKdm2*^*2*^ stable homozygous lines were established with standard crossing protocol.

To further validate these mutant alleles, we extracted the genomic DNA from 10 homozygous *dKdm2*^*1*^and *dKdm2*^*2*^ mutant larvae using DNAzol (Invitrogen, 10503-027). PCR was performed with Taq polymerase (Invitrogen) for 35 cycles (elongation for 5.5min and annealing at 55°C), as described previously (Zheng *et al.* 2014). The following primers were used: *dKdm2-3F*: 5’-CGGTTGTAGCCGTTAGGAAA; *CRISPR dKdm2-5.1*: 5’-CACGCAAGAATGCAGAGGTA; and *CRISPR dKdm2-3.1*: 5’-TTTGCATTGCGTGTGGTTAT. The PCR products were further validated by sequencing.

### Generation of the *dKdm2*^*+*^*-eGFP* line

To tag the endogenous *dKdm2* locus with eGFP, we used the germ-line specific CRISPR/Cas9 system (Ren *et al.* 2013; Ren *et al.* 2014). One sgRNAs were designed to target 3’UTR of *dKdm2* (CTGGCAGCTGGAGGAGGGCC) and cloned into U6b-sgRNA plasmid. The *dKdm2*^*+*^*-eGFP* donor construct was based on the pBluescript plasmid. The left homologous arm of *dKdm2* was amplified from genomic extract with *dKdm2*-*left-F* primer (5’-TCTCTCAGTTGGGGAAGCTTCCACTAATCAGTGGAGTGGCAGTGGC) and *dKdm2-left-R* primer (5’-TCCGCTTCCTCCTCCTCCGCTTCCACCTCCTCCGTCGTGGTGCCAGCTTCGGT). The right homologous arm was amplified with *dKdm2-right-F* primer (5’-GCGACTGGCAGCTGGAGGAGGGC) and *dKdm2-right-R* primer (5’-TCTCTCAGTTGGGGAAGCTTAAGATGCAGGACGGTCGCAAGACG). The gene encoding the *eGFP* was amplified with *eGFP-F* primer (5’-GGAGGAGGAGGAAGCGGAGGAGGAGGTAGCGTGAGCAAGGGCGAGGAGCTG) and *eGFP-R* primer (5’-CTCCTCCAGCTGCCAGTCGCTTACTTGTACAGCTCGTCCATGCCGAG). GS linker was synthesized in *eGFP-F* and *dKdm2-left-R* primers. Fragments were stitched together using the sequence- and ligation-independent cloning (Jeong *et al.* 2012). All PCR fragments were amplified with KOD DNA polymerase (TOYOBO). The donor construct was confirmed by sequencing with *eGFP-seq-R* primer (5’-ACCCCGGTGAACAGCTCCT) and *dKDM2-seq-F* primer (5’-GCGGCGACTCCAAAGTAGGAAA). To generate *dKdm2-eEFP* flies, sgRNA plasmids and donor were co-injected into nos-Cas9 *Drosophila* embryos. All G0 adult flies that developed from injected embryos were crossed with the “*y*^*1*^ *sc*^*1*^ *v*^*1*^; *+; Dr*^*1*^ *e*^*1*^*/TM3 Sb*^*1*^” line, knock-in events were screened by PCR of F1 adults and validated by sequencing. The *dKdm2-eGFP* stable homozygous lines were established using standard fly genetics.

### Characterization of the *dKdm2* alleles using Western blot and qRT-PCR

The Western blot analyses of the *dKdm2* alleles were performed as described (Zheng *et al.* 2014). A Li-Cor Odyssey Infrared Imaging system was used to quantify the Western blots shown in Fig. 2 and Fig. 3. The primary antibodies used are described previously (Zheng *et al.* 2014), and the following secondary antibodies from Li-Cor Biosciences are used in these analyses: IRDye 800CW Goat anti-Rabbit IgG (926-32211) and IRDye 680 Goat anti-Mouse IgG (926-32220). At least three independent biological repeats were quantified in each experiment. The primers and conditions for qRT-PCR assays were the same as described previously (Zheng *et al.* 2014). Statistical significance was determined using one-tailed Student’s t-test.

**Fig. 1.**
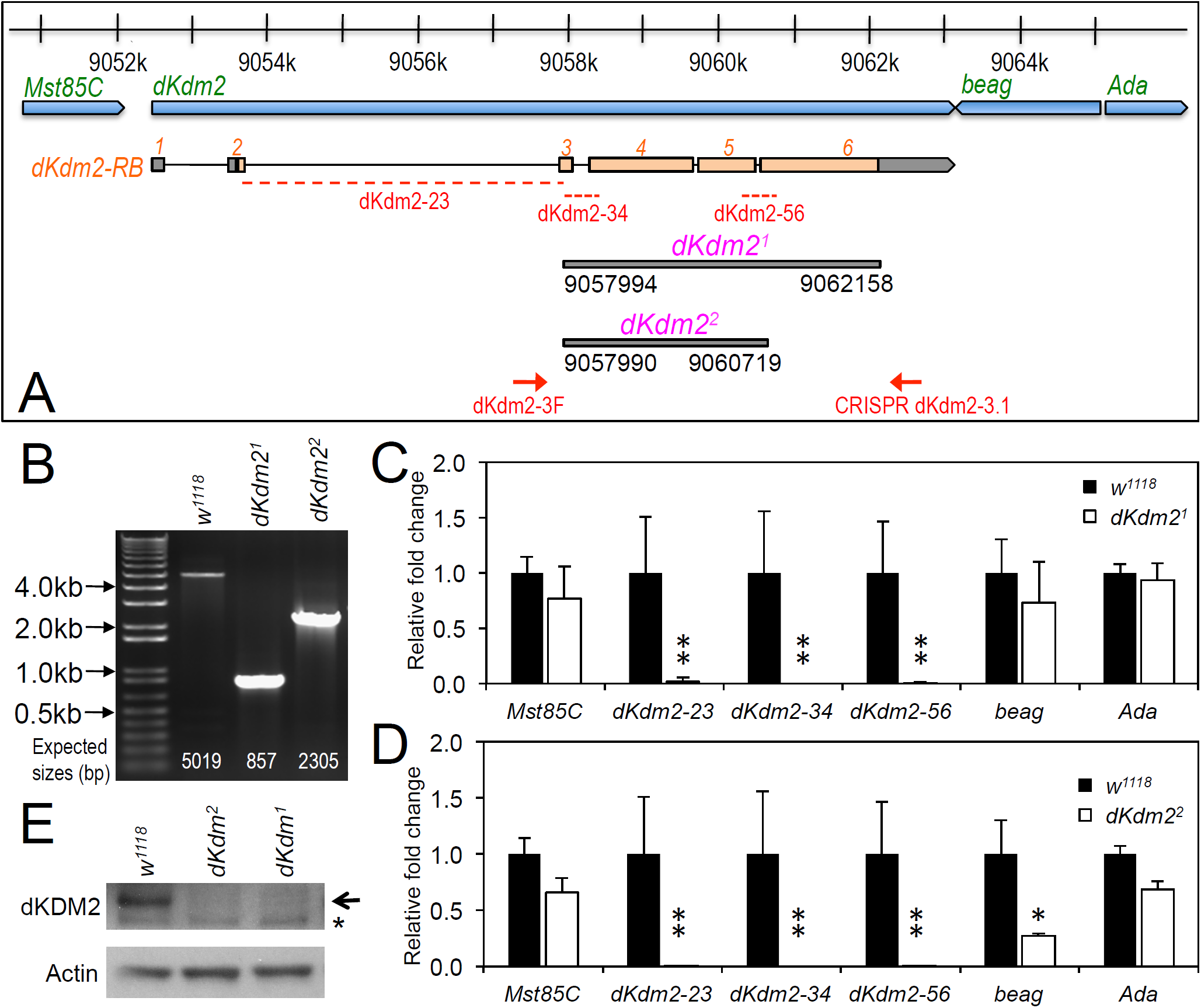
Generation and characterization of the *dKdm2* null alleles. (A) Schematic drawing of the *dKdm2* locus and its neighboring genes, note the regions that are deleted in *dKdm2*^*1*^ and *dKdm2*^*2*^ and the corresponding breakpoints. (B) Validation of the *dKdm2* alleles using PCR and genomic DNA. The positions of the primers (“dKdm2-3F” and “CRISPR dKdm2-3.1”) are shown in (A). (C, D) The mRNA levels of *dKdm2* and its neighbor genes *dKdm2, Mst85C, beag* and *Ada* in the third instar larvae are measured by qRT-PCR assay. The positions of the primers for three regions within the *dKdm2* locus are shown in (A). (E) The protein levels of dKDM2 in the third instar (L3) wandering larvae were analyzed by Western blot. The non-specific bands are marked with ‘*’, and anti-actin was used as the control.

**Fig. 2.**
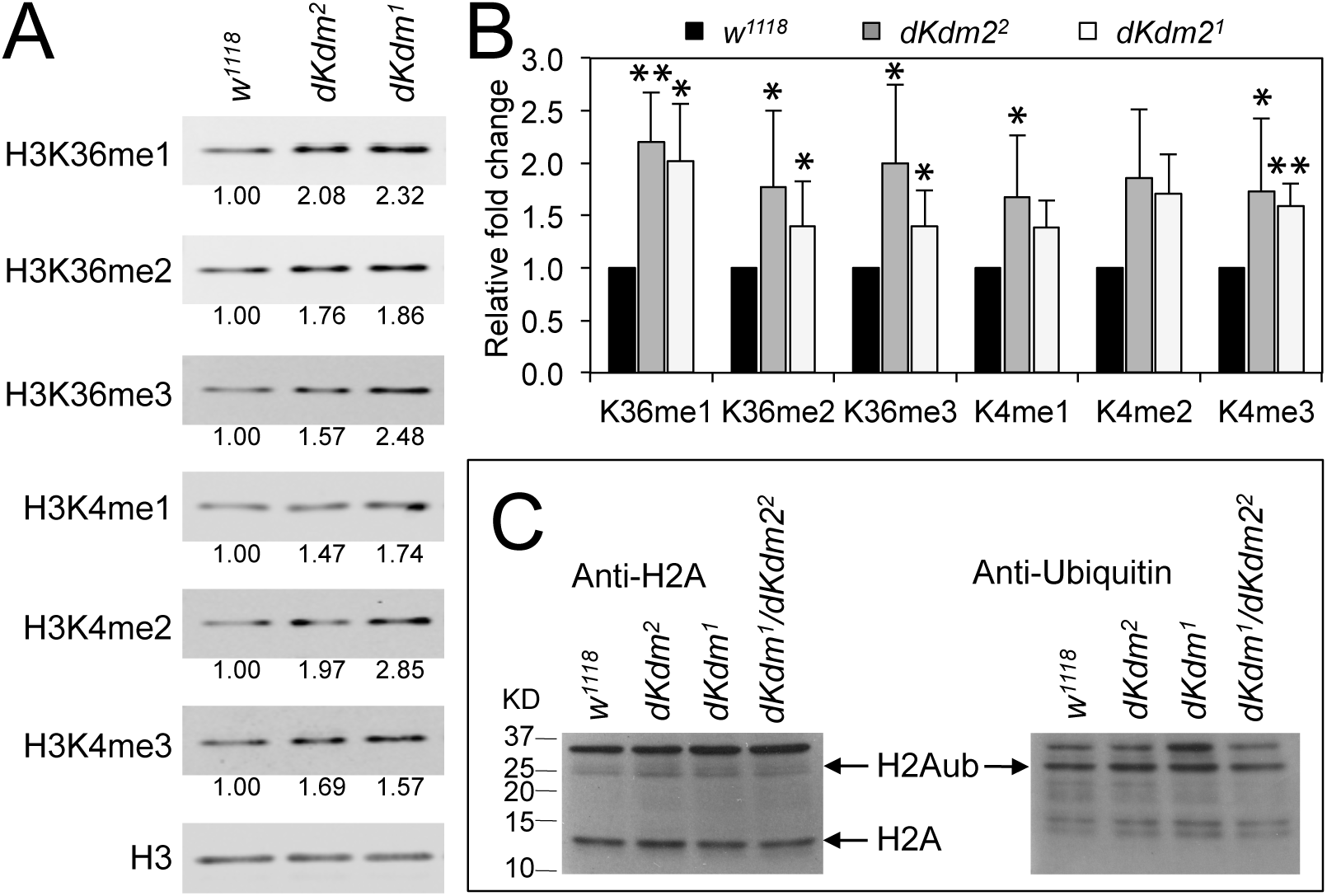
Levels of histone methylation and ubiquitination in the *dKdm2* mutants. (A) The levels of H3K36me1/2/3 and H3K4me1/2/3 in the *dKdm2* mutants at the L3 wandering stage. The results of Western blot (A) from three independent biological repeats were quantified using a Li**-**Cor Odyssey Infrared Imaging system (B). * p < 0.05; ** p < 0.01 based on Student’s *t*-tests. (C) The levels of histone H2A and ubiquitinated H2A (H2Aub) the *dKdm2* mutants at the L3 wandering stage.

**Fig. 3.**
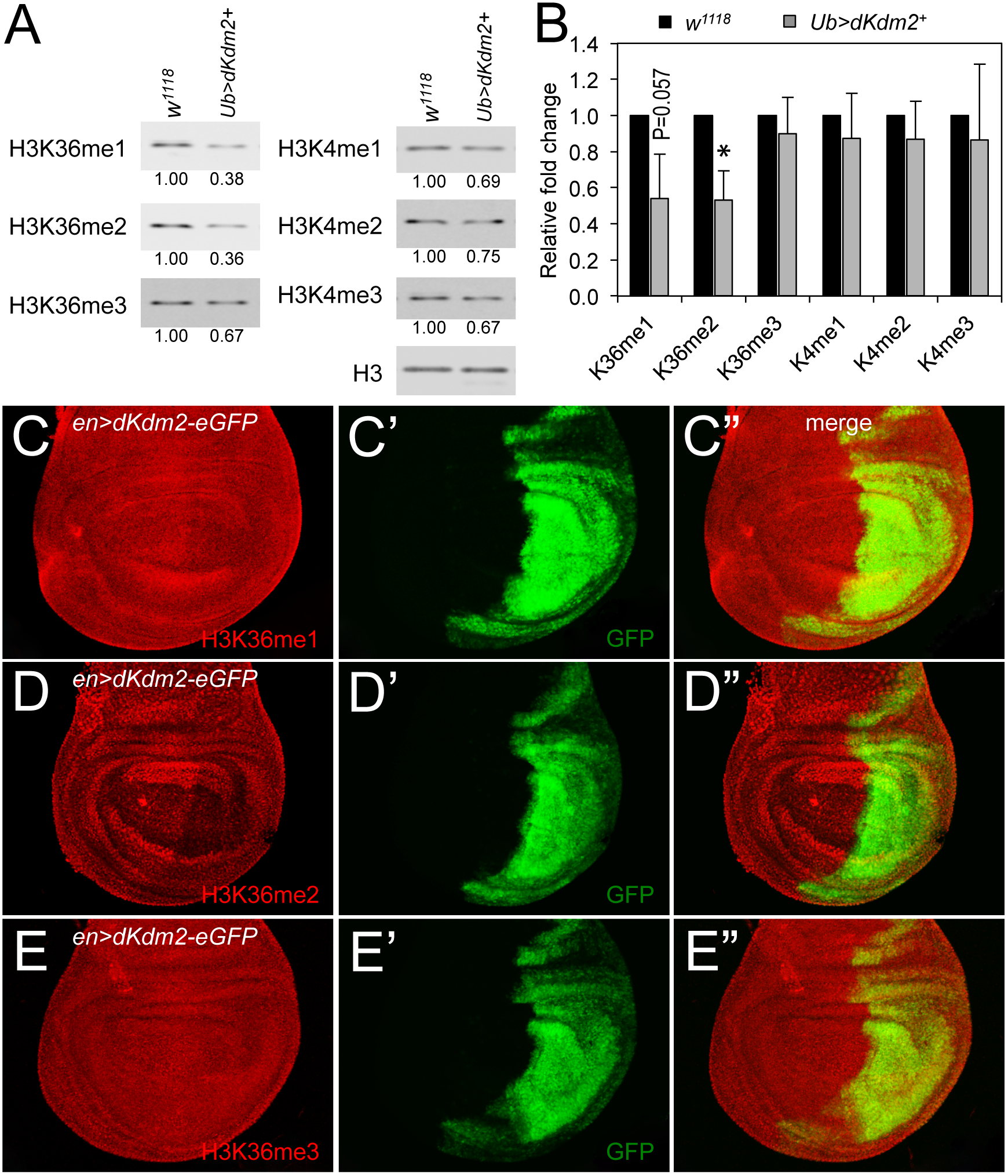
Effects of dKDM2 over-expression on methylation levels of H3K4 and H3K36. (A) The levels of H3K36me1/2/3 and H3K4me1/2/3 in dKdm2-overexpressed larvae at the L3 wandering stage. The detailed genotype for “*Ub>dKdm2*^*+*^” line is “*w*^*1118*^; *Ubi-Gal4/UAS-dKdm2-eGFP; +*”. (B) Quantification of the results of Western blots from three independent biological repeats using a Li**-**Cor Odyssey Infrared Imaging system. (C-E) Immunostaining of wing discs with wild-type *dKdm2* overexpressed in the posterior half of the discs (genotype: *en-Gal4/UAS-dKdm2-eGFP*) with antibodies against H3K36me1 (C), H3K36me2 (D), and H3K36me3 (E). Images in C’-E’ show the GFP signals, and C”-E” are merged images.

### Immunostaining procedure of wing discs

Wing discs from the third-instar wandering larvae were dissected in phosphate buffered saline (PBS), and then fixed in 5% formaldehyde in PBS for 10 min at room temperature. After rinsing with PBT (PBS with 0.2% Triton X-100), the discs were blocked with PBTB (PBT with 0.2% BSA and 5% normal goat serum) for 1h at room temperature with rocking. The discs were then incubated with the following primary antibodies: anti-H3K4me2 was purchased from Cell Signaling Technology (Danvers, 9725; 1:500 diluted in PBTB); while anti-H3K4me1 (ab8895; 1:2500), anti-H3K4me3 (ab8580; 1:500), anti-H3K36me1 (ab9048; 1:500), anti-H3K36me2 (ab9049; 1:1000), and anti-H3K36me3 (ab9050; 1:500) were purchased from Abcam. After rinsing, the discs were then incubated with the secondary antibody (1:500 in PBTB) for 1 hr at room temperature on a nutator. Finally, the discs were rinsed with PBT and mounted with VECTASHIELD. Confocal images were collected using a Nikon Ti Eclipse microscope, and then processed using the Adobe Photoshop CS6 software.

### Analyses of the circadian locomotor behaviors

2-5 days old male flies were collected and loaded into *Drosophila* Activity Monitor (DAM, TriKinetics). Flies were entrained to 12 h light: 12 h dark cycles (LD) at 25°C for 4-5 days and then were released into constant darkness (DD) for 7 days. Locomotor activity rhythms were analyzed using FaasX software. Average actograms were generated by MATLAB with Griffith sleep analysis toolbox, kindly shared from Fang Guo in Michael Rosbash lab.

### Whole-mount brain immunohistochemistry

Adult flies were entrained in LD for 4-5 days, and then were collected at the indicated circadian time (CT) on the first day of DD. Flies were immediately fixed in 4% paraformaldehyde in PBS at room temperature for 1 hour and further were dissected in PBS containing 0.1% Triton-X100 (PBST). After three times washes using PBST, fly brains were blocked in PBST with 10% normal goat serum at room temperature for 1 hour. Next, they were incubated at 4°C overnight with primary antibody: rabbit anti-PER (1:5000), guinea pig anti-CLK (1:2500), and mouse anti-PDF (C7, Developmental Studies Hybridoma Bank, 1:400). Secondary antibodies were diluted at 1:200 concentration with respect to anti-mouse 488, anti-rabbit 594, and anti-guinea pig Cy3 (Jackson Immuno Research). For dKDM2-EGFP expression, brains were dissected and staining with an anti-GFP antibody (Invitrogen, A6455; 1:200) and mouse anti-PDF (1:400). Brains were imaged using Leica TCS SP8 confocal microscope and the images were quantified using the NIH ImageJ (https://imagej.nih.gov/ij/). Statistical significance were determined by two-tailed Student’s t-test at P < 0.05.

### Data Availability Statement

All strains and plasmids are available upon request. The authors affirm that all data necessary for confirming the conclusions of the article are present within the article, figures, and Tables.

## Results

### Generation of the *dKdm2* null alleles using the CRISPR/Cas9 technique

To determine the role of dKDM2 in *Drosophila* development, we generated two mutant alleles that only delete the coding region of *dKdm2*, designated as *dKdm2*^*1*^ and *dKdm2*^*2*^ (Fig. 1A), using the CRISPR/Cas9 technique (Ren *et al.* 2014; Bier *et al.* 2018). These two alleles are null alleles of *dKdm2* gene based on the following evidences. First, by performing PCR using the genomic DNA from the third instar larvae of these two mutant lines, we observed shorter PCR products in the mutants compared to the control as expected (Fig. 1B). Second, sequencing the PCR products has revealed that *dKdm2*^*1*^ deletes the region 9057994∼9062158 of the third chromosome, and *dKdm2*^*2*^ deletes the region 9057990∼9060719 (Fig. 1A). Third, by analyzing the mRNA levels of *dKdm2* using qRT-PCR assay, the transcripts of *dKdm2* are undetectable, in contrast to *w*^*1118*^ control and the neighboring genes (Figs. 1C and 1D). Furthermore, the dKDM2 protein is not detectable from the mutant larvae as assayed by Western blot (Fig. 1E). These analyses demonstrate that *dKdm2*^*1*^ and *dKdm2*^*2*^ disrupt the *dKdm2* locus. For unknown reasons, the mRNA level of *beag*, a neighboring gene of *dKdm2*, is also reduced in the *dKdm2*^*2*^ mutants, as assayed by multiple repeats. Nevertheless, our analyses with both these alleles did not reveal any difference between them (see below). These results show that both *dKdm2*^*1*^ and *dKdm2*^*2*^ are null alleles.

Importantly, the homozygous mutants of *dKdm2*^*1*^ and *dKdm2*^*2*^, as well as the transheterozygous animals (genotype: *w*^*1118*^; *+; dKdm2*^*1*^*/dKdm2*^*2*^), are fully viable and fertile, without any developmental defects. We also did not observe any differences between these mutants and the control on their life spans and fertilities (data not shown). In fact, the homozygous mutants of *dKdm2*^*1*^ and *dKdm2*^*2*^ can be maintained as stable and healthy stocks for many generations. Taken together, these results demonstrate that *dKdm2* is not required for normal development. This conclusion is consistent to our previous work (Zheng *et al.* 2014), as well as a recent report of a null *dKdm2* allele independently generated by an ends-out gene targeting approach (Shalaby *et al.* 2017).

### Effects of loss of *dKdm2* on histone methylation and ubiquitination

Previously, we reported that methylation on H3K36 and H3K4 are only weakly affected in the deficiency or transposon insertion lines in the *dKdm2* locus (Zheng *et al.* 2014). With these two new *dKdm2* null alleles, we re-examined whether on the methylation status on H3K36 and H3K4 are affected in the *dKdm2* null mutants using quantitative Western blot analysis. As shown in Fig. 2A, the levels of H3K36me1/2/3 and H3K4me1/2/3 are elevated in the *dKdm2*^*1*^ or *dKdm2*^*2*^ homozygous mutants at the L3 wandering stage. Quantification of the Western blots from three independent experiments show that the increases on H3K36me1/2/3 and H3K4me1/3 are statistically significant (Fig. 2B). These results are consistent to the previous reports that dKDM2 may be involved in regulation the demethylation of H3K4 and H3K36 (Lagarou *et al.* 2008; Kavi and Birchler 2009; Zheng *et al.* 2014).

Depletion of dKDM2 in S2 cells was shown to significantly increased H2AK118ub (Lagarou *et al.* 2008). It has been reported that the mouse KDM2B is required for H2AK119ub (Wu *et al.* 2013). However, a recent report has identified the RING1-L(3)73Ah complex, which does not contain dKDM2, as the major regulator of the H2AK118ub in *Drosophila* (Kahn *et al.* 2016). Thus we asked whether the levels of H2A ubiquitination is altered in the *dKdm2* null mutant larvae. As shown in Fig. 2C, the ubiquitin levels on H2A are not affected in the *dKdm2*^*1*^, *dKdm2*^*2*^, or *dKdm2*^*1*^/*dKdm2*^*2*^ transheterozygous mutant larvae at the wandering stage. This observation does not support a role of dKDM2 in regulating H2A ubiquitination. Taken together, these results suggest that the major role of dKDM2 is to regulate the methylation states on H3K36 and H3K4.

To further analyze the role of dKDM2 on histone modifications, we asked whether over-expression of wild-type dKDM2 could have opposite effects on H3K36 and H3K4 methylation status. To test this, we generated *UAS-dKdm2* transgenic lines, which allow us to overexpress wild-type *dKdm2* in a tissue-specific or developmental stage-specific manner (Fig. S1). We ectopically expressed dKDM2 in all cells using the *ubi-Gal4* driver and then analyzed histone methylation in L3 wandering larvae by Western blot. As shown in Fig. 3A, overexpression dKDM2 reduced levels of H3K36me1 (slightly) and H3K36me2 (significantly), but no obvious effects were observed on levels of H3K36me3 and H3K4me1/2/3. This notion is confirmed by the quantitative Western blot analyses based on three independent repeats (Fig. 3B). These data suggest that gain of dKDM2 is sufficient to reduce the H3K36me1 and H3K36me2 levels.

To complement these biochemical analyses, we performed immunostaining using the wing discs in which wild-type dKDM2 is ectopically expressed in the posterior compartment of the wing discs using the *en-Gal4* driver. As shown in Fig. 3C, the levels of H3K36me1 is barely reduced in the posterior part of the wing disc, shown by simultaneously expressed GFP (Fig. 3C’). In contrast, the levels of H3K36me2 is clearly reduced in cells with overexpression of wild-type dKDM2 (Figs. 3D and 3D’), but no regional difference was observed with H3K36me3 (Fig. 3E) or H3K4me1/2/3 (Suppl. Fig. S2). These observations based on the immunostaining approach are consistent to the biochemical analysis (Figs. 3A and 3B). Taken together, these results suggest that H3K36me2 is the major target of dKDM2 during the larval stage.

### *dKdm2* mutants are defective in circadian rhythms

Considering the high evolutionary conservation of the KDM2 family of enzymes and the fundamental roles of H3K36 methylation in transcription, it is puzzling that *dKdm2* null mutants do not display any developmental defects (Zheng *et al.* 2014). One clue may be obtained by analyzing the expression of the endogenous dKDM2 proteins, but this effort is hampered by the lack of the specific antibodies suitable for immunostaining of dKDM2. After our initial report on the 10 transposon insertion lines within the *dKdm2* locus, we learned that another insertion line *dKdm2*^*36Y*^ (*w*; p[GawB]dKdm2*^*36Y*^) in the *dKdm2/CG11033* locus became available. This Gal4 enhancer trap line, also known as *36Y-Gal4*, is caused by the insertion of the *p[GawB]* transposon in the first intron of the *dKdm2* locus (Taghert *et al.* 2001). At the adult stage, this Gal4 line is expressed in the central brain, optic lobes, ring gland, salivary glands, fat body, epidermis, and hindgut (Taghert *et al.* 2001). *36Y-Gal4* is expressed in peptidergic neurons of the CNS and interestingly, *36Y-Gal4* driven expression of the amidating enzyme PHM affects the circadian rhythms in *Drosophila* (Taghert *et al.* 2001). In addition, using cultured human osteosarcoma U2OS cell, KDM2A/FBXL11 has been identified from a screen for F-box proteins required for normal circadian rhythms assayed by the expression of a clock reporter (Reischl and Kramer 2015). Therefore, we asked whether dKDM2 is involved in regulating circadian locomotor rhythms.

There are four large ventral lateral neurons (lLNvs) and four small ventral lateral neurons (sLNvs) that express the neuropeptide pigment dispersing factor (PDF) in each fly brain hemisphere (Nitabach and Taghert 2008). The PDF positive sLNvs are the master pacemaker neurons that control the circadian rhythms in constant darkness (DD) (Renn *et al.* 1999). Although *36Y-Gal4* is expressed in the lLNvs and sLNvs, it remains unknown if the expression pattern of precisely represent the expression pattern of the *dKdm2* gene (Taghert *et al.* 2001). Thus to test whether the endogenous dKDM2 protein is expressed in adult brains, we inserted an eGFP tag in the C-terminus of the endogenous *dKdm2* locus using the CRISPR-Cas9 technique (see Materials and Methods). The *dKdm2-eGFP* flies are fully viable, and we have observed that dKDM2 is expressed adult brains (Fig. S2A), including the sLNvs (Fig. S2B) and lLNvs (Fig. S2C). The dKDM2 protein is predominantly localized in neucleus (Fig. S2), consistant with its function in regulation of histone demethylations and our previous report (Zheng *et al.* 2014).

Encourgaed by these observations, we analyzed the circadian behaviors of the *dKdm2*^*1*^ or *dKdm2*^*2*^ homozygous mutants. They displayed defects of circadian locomotor rhythms under DD. For *dKdm2*^*1*^, only ∼37% of the flies were rhythmic, but circadian period was shortened for ∼1 hour (Fig. 4A, Table 1). Similar phenotypes were also observed in *dKdm2*^*2*^ mutants, as well as the transheterozygous animals (Fig. 4A, 4B). When we further checked the locomotor behavior under light/dark (LD) cycle, we identified that the morning anticipatory behavior peak was also blunted in *dKdm2*^*1*^ or *dKdm2*^*2*^ mutants (Fig. 4C). Consistent with our *dKdm2* mutants phenotype, when we overexpressed wild-type dKDM2 in all circadian neurons using the *tim-Gal4* driver, the period was increased for ∼1 hour compared to the controls (Fig. 4B, 4D, Table 1). Given that *36Y-Gal4* has weak expression in the sLNvs, we futher tested the circadian behavior of flies with dKDM2 overexpression in sLNvs using the *pdf-Gal4* driver. Similar lengthened circadian period phenotype was observed (Fig. 4B). However, a weak but significant period lengthening was also identified when we overexpressed wild-type dKDM2 in all circadian tissues except the sLNvs (TG4; PG80, Fig. 4B, Table 1). These results suggest that dKDM2 regulates circadian period in sLNvs, but it may also fine-tune the rhythms in PDF negative circadian neurons.

**Table 1.**
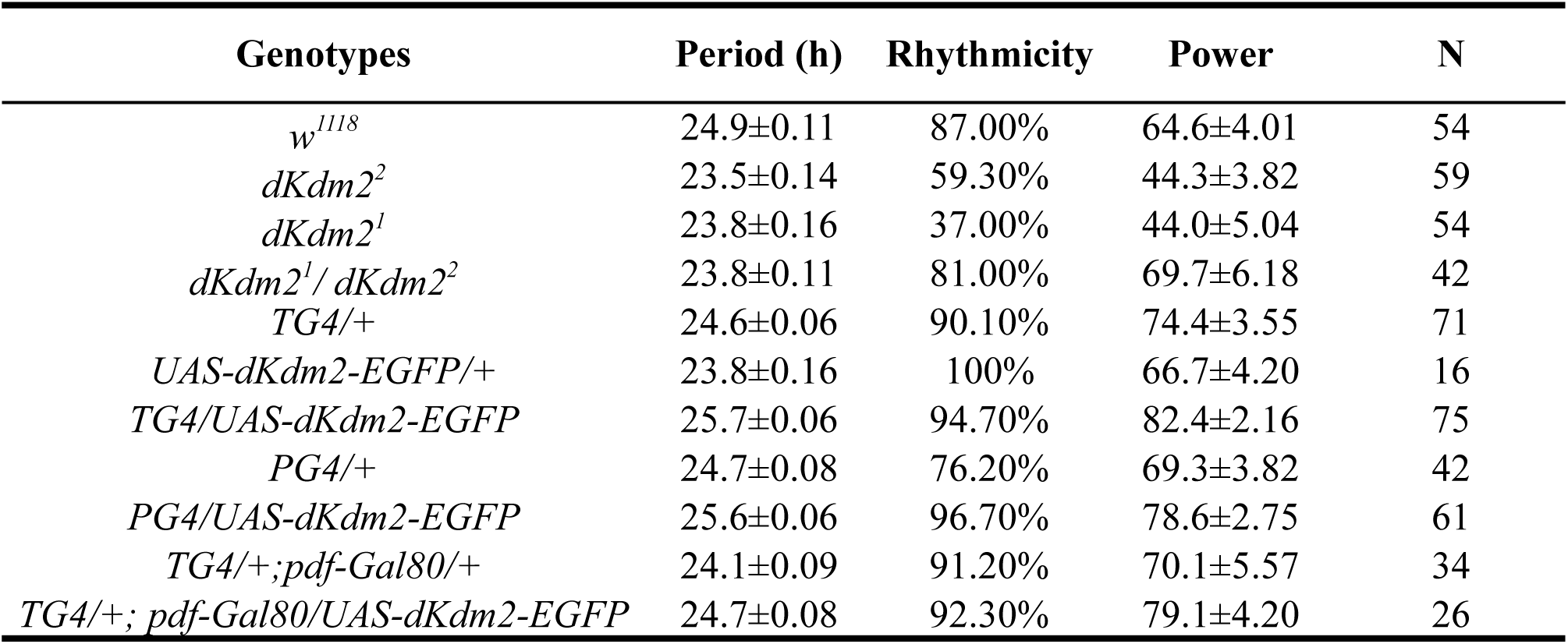
Effects of dKDM2 mutation or overexpression on circadian behaviors.

| **Genotypes** | **Period (h)** | **Rhythmicity** | **Power** | **N** |
| --- | --- | --- | --- | --- |
| *w<sup>1118</sup>* | 24.9±0.11 | 87.00% | 64.6±4.01 | 54 |
| *dKdm2<sup>2</sup>* | 23.5±0.14 | 59.30% | 44.3±3.82 | 59 |
| *dKdm2<sup>1</sup>* | 23.8±0.16 | 37.00% | 44.0±5.04 | 54 |
| *dKdm2<sup>1</sup>/dKdm2<sup>2</sup>* | 23.8±0.11 | 81.00% | 69.7±6.18 | 42 |
| *TG4/+* | 24.6±0.06 | 90.10% | 74.4±3.55 | 71 |
| *UAS-dKdm2-EGFP/+* | 23.8±0.16 | 100% | 66.7±4.20 | 16 |
| *TG4/UAS-dKdm2-EGFP* | 25.7±0.06 | 94.70% | 82.4±2.16 | 75 |
| *PG4/+* | 24.7±0.08 | 76.20% | 69.3±3.82 | 42 |
| *PG4/UAS-dKdm2-EGFP* | 25.6±0.06 | 96.70% | 78.6±2.75 | 61 |
| *TG4/+;pdf-Gal80/+* | 24.1±0.09 | 91.20% | 70.1±5.57 | 34 |
| *TG4/+;pdf-Gal80/UAS-dKdm2-EGFP* | 24.7±0.08 | 92.30% | 79.1±4.20 | 26 |

**Fig. 4.**
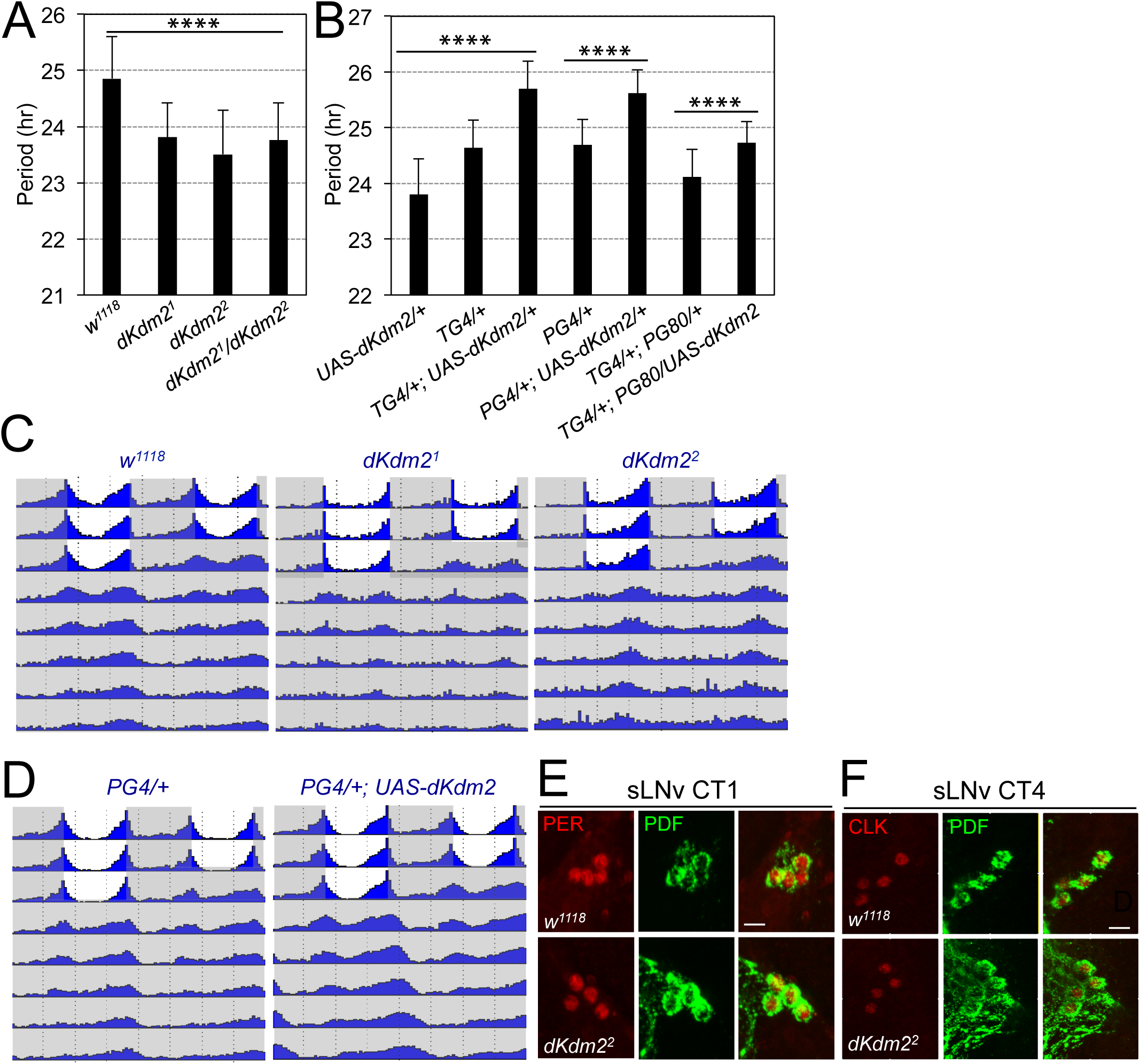
dKDM2 is required for circadian behavior rhythms. (A) Period of circadian locomotor rhythms of the *dKdm2* mutants. (B) Effects of *dKdm2* overexpression on the period of circadian locomotor rhythms. Error bars indicate S.D.; the detailed genotype of “*TG4/+*” is “*tim-GAL4, UAS-dicer2/+*”; “*PG4/+*” is “*Pdf-GAL4, UAS-dicer2/+*”; and “*PG80*” is “*Pdf-GAL80*”. Each genotype is compared to its driver or UAS line as the controls. For the data shown in (A) and first three columns in (B), ****p < 0.0001, as determined by using Tukey’s multiple comparison test after one-way analysis of variance. For the rest of the data shown in (B), ****p < 0.0001, based on *t*-test. Number of animals analyzed is included in Table 1. (C, D) Locomotor behavior under light dark (LD) cycle and constant darkness (DD). Representative double plotted actograms showing average activity of flies during 3 days of LD and 5 days of DD for controls, *dKdm2* mutants (C) and *dKdm2* overexpression (D) in PDF positive sLNvs using the *Pdf-GAL4* driver. The light phase is shown in white color, and the dark phase is shown in gray. (E, F) Representative confocal images showing the effects of *dKdm2* mutation on the levels of PER (E) and CLK (F) in sLNvs. Flies were entrained under 3 days of LD, and brains were dissected at CT0 (CT= circadian time) in the first day of DD. Fly brains were dissected at CT4. Scale bars in E/F: 10 µm.

The core of circadian clocks contains transcriptional translational feedback loops that are highly conserved among species (Hardin and Panda 2013). In *Drosophila*, a transactivator complex formed by CLOCK (CLK) and CYCLE (CYC) generates rhythmic transcription of hundreds of genes, including some transcriptional repressors; PERIOD (PER) is the major repressor, which accumulates in the cytoplasm, undergoes a series of post-translational modifications, and entry into the nuclear to block its own transcription (Hardin and Panda 2013). To understand how dKDM2 might affect circadian behavior, we compared protein abundance of PER and CLK at the time point of their peak expression under constant darkness, circadian time 1 (CT1) for PER (circadian time) and CT4 for CLK, respectively (Fig 4C). No obvious changes of PER (Fig. 4E, quantified in Fig. S3A) and CLK (Fig. 4F, S3B) levels were observed in either sLNv, or lLNvs (Fig. S3C/C’, Fig. S3D/D’). These observations indicate that dKDM2 does not affect the abundance of the core pacemaker proteins. Taken together, these results show that dKDM2 is involved in regulating circadian rhythms, and varying its dosage affects the circadian rhythmicity and period.

## Discussion

Characterization of the phenotypes of different mutant alleles that disrupt the functions of a specific gene is essential to understand how its product functions during the development of an organism. In this study, we report the generation and characterization of two null alleles of the *dKdm2* gene that specifically disrupt *dKdm2* locus but not its neighboring genes. The homozygous mutant animals are fully viable and fertile, but are defective in circadian rhythms. In addition, the null mutant larvae display mild effects on methylation states on H3K36 and H3K4 during the larval stage, but no effect on the levels of H2Aub was observed. Moreover, overexpression of wild-type dKDM2 in larvae and wing discs reduced the levels of H3K36me2, but not the levels of H3K36me3 and H3K4me1/2/3. Furthermore, *dKdm2* null mutants display a shortened circadian period, while overexpression of dKDM2 increased the circadian period. These results show that dKDM2 is involved in regulating circadian clocks but does not play essential roles in normal development, and the major targets of dKDM2 *in vivo* are H3K36me2. These results also identify the major function of dKDM2 as an epigenetic regulator of circadian rhythms.

### dKDM2 is not an essential gene for viability in *Drosophila*

dKDM2 was initially identified as a subunit of the dRAF complex that couples H2Aub to demethylation of H3K36me2 during Polycomb group silencing (Lagarou *et al.* 2008). Subsequently, dKDM2 was shown to demethylate H3K4me3 (Kavi and Birchler 2009). Given the fundamental roles of Polycomb complexes in regulating the body plan in animals and the methylation states of H3K4 and H3K36 in regulating transcription activation and elongation, dKDM2 was expected to be critical for development. However, developmental genetic analyses of multiple *dKdm2* mutant alleles in *Drosophila* suggest that *dKdm2* is not required for normal development (Zheng *et al.* 2014; Shalaby *et al.* 2017), and this work using two null alleles of *dKdm2* has further validated this conclusion.

It has been reported that *dKdm2* genetically interacts with *Pc*^*1*^, *Pc*^*3*^, *trx*^*1*^ and *ash1*^*10*^ (Lagarou *et al.* 2008). Three mutant alleles of *dKdm2* were used in these genetic tests, including *dKdm2*^*DG12810*^, *dKdm2*^*EY01336*^, and *dKdm2*^*KG04325*^; of these, *dKdm2*^*DG12810*^ and *dKdm2*^*KG04325*^ were reported to be homozygous lethal alleles, while *dKdm2*^*EY01336*^ is a homozygous viable allele (Lagarou *et al.* 2008). In another study, *dKdm2*^*DG12810*^ was shown to genetically interact with the KDM5 homolog *little imaginal disc* (*lid*) (Li *et al.* 2010). However, these alleles were not validated using molecular and biochemical analyses before their usage. In our previous analyses, the *dKdm2*^*EY01336*^ and *dKdm2*^*KG04325*^ homozygotes are viable; for both of these alleles, the transposons inserted in the second intron of *dKdm2* gene have no effects on either the mRNA or protein levels of dKDM2 (Zheng *et al.* 2014). After outcrossing the *dKdm2*^*EP3093*^, *dKdm2*^*f02828*^, and *dKdm2*^*d00170*^ mutants, which were initially homozygous lethal, with the wild-type flies for four generations, we have observed that the homozygous of these alleles are fully viable, thus these original alleles carry second site lethal mutations (Zheng *et al.* 2014). However, using the same outcrossing strategy, the *dKdm2*^*DG12810*^ homozygous mutants are still lethal at the third instar larval and pupal stages, yet the transheterozygous of this allele with other strong *dKdm2* alleles and deletion lines are fully viable (Zheng *et al.* 2014). These observations suggest that the second site lethal mutation(s) in the *dKdm2*^*DG12810*^ allele is close to the *dKdm2* locus thus cannot be easily removed. Given that the existence of second-site mutations can complicate the interpretation of genetic analyses, it is important to validate mutant alleles, especially when they are being used for the first time.

### Role of dKdm2 in demethylating histone H3

dKDM2 protein has several conserved protein domains, including the JmjC, CXXC-type zinc finger, F-box, and Amn1 (Antagonist of mitotic exit network protein 1) domains (Zheng *et al.* 2014; Holowatyj *et al.* 2015). Of these domains, the JmjC domain defines its enzymatic activity as a demethylase. Considerable efforts have been invested into elucidating the exact histone markers of KDM2 proteins *in vitro* and *in vivo.*

Based on *in vitro* demethylation assay using modified histone H3 peptides as the substrates followed with mass spectrometry analysis, it has been shown that KDM2A preferentially demethylate H3K36me2, but not H3K36me3, which was validated by forced expression of KDM2A in 293T cells (Tsukada *et al.* 2006). Similar approach has been used to show that KDM2A only demethylates H3K36me1 and H3K36me2, but not other methylated peptides such as H3K36me3, H3K4me1/2/3, H3K9me1/2/3, and H3K27me1/2/3 (Williams *et al.* 2014). Moreover, structural analyses demonstrate that KDM2A specifically demethylates H3K36me2 (Cheng *et al.* 2014). In contrast, the results from analyses using *in vivo* samples are a less clear, largely dues to different experiments systems and approaches. Both KDM2A and KDM2B can specifically demethylate H3K36me2 (He *et al.* 2008; Kottakis *et al.* 2011; Liang *et al.* 2012). Overexpression of KDM2A in HeLa cells decreased the levels of H3K36me2; while ectopic expression of KDM2B significantly reduced the H2K4me3 level but has no effects on the levels of H3K4me2, H3K9me3, H3K27me3, H3K36me2, and H3K36me3 (Frescas *et al.* 2007). However, expression of KDM2B in HEK293 and HeLa cells only reduced the levels of H3K36me2, but not H3K4me3 (He *et al.* 2008). Expression of KDM2B in mouse embryonic fibroblasts also significantly reduced the levels of H3K36me2 (Liang *et al.* 2012). Furthermore, the JmjC domain of KDM2B tagged with GST can demethylate H3K4me3 but not H3K36me2 in *in vitro* histone demethylation assay (Janzer *et al.* 2012). These studies suggest that H3K36me2 is the major target for KDM2A and KDM2B in mammalian cells, and KDM2B can also demethylate H3K4me3.

Depletion of dKDM2 in cultured *Drosophila* S2 cells increases the H3K36me2 level but not the levels of H3K36me1/3 or H3K4me3 (Lagarou *et al.* 2008), yet knocking down of *dKdm2* in larvae increased the levels of H3K4me3 but not H3K36me2 (Kavi and Birchler 2009). In our hands, we observed mild elevation of the H3K36me1/2/3 levels in transheterozygous larvae of deletions lines uncovering the *dKdm2* locus, and we could not detect changes on the global levels of H3K36me1/2/3, H3K4me1/2/3, H3K9me2, and H3K27me2/3 in S2 cells (Zheng *et al.* 2014). Using a more quantitative method for Western blot, we observed a mild increase in the levels of H3k36me1/2/3 and H3K4me1/2/3 in *dKdm2* null mutant larvae (Fig. 2). Interestingly, however, when wild-type dKDM2 is overexpressed in larvae, we can detect considerable reduction in the level of H3K36me2 and to a less extent, the H3K36me1 level, but not the levels of H3K36me3 and H3K4me1/2/3 (Fig. 3A/3B). These observations are validated by immunostaining of these epigenetic markers in wing discs, where the cells in the anterior compartment serve as the internal control (Fig. 3C-3E). Taken together, these different results are likely due to different experimental approaches; considering all the observations summarized above and the intrinsic strengths and limitations of different methods, it appears that most data support H3K36me2 as the major target of the dKDM2. Perhaps, instead of analyzing the global effects of dKDM2 on epigenetic markers, analyzing the levels of these modifications on the specific promoters may provide better resolution in the future.

### Role of dKDM2 in regulating H2A ubiquitination

The histone H2A lysine 118 (H2AK118) in *Drosophila* is mono-ubiquitinated by Sce (Sex combs extra, or dRING), and its vertebrate orthologs Ring1 and Ring2/Ring1B act as the E3 mono-ubiquitin ligase for the corresponding H2A at K119 (de Napoles *et al.* 2004; Wang *et al.* 2004; Lagarou *et al.* 2008; Gutierrez *et al.* 2012). The dimerization between RING1 and a member of the family of Polycomb group ring finger (PCGF) proteins is necessary for the H2A ubiquitinase activity of RING1 (Wang *et al.* 2004; Cao *et al.* 2005). In *Drosophila*, the PCGF proteins include Posterior sex combs (Psc), Suppressor of zeste 2 (Su(z)2), and L(3)73Ah (Dorafshan *et al.* 2017). The dKDM2 has been identified as a key subunit of the dRAF complex, composed of dRING, Psc, dKDM2, RAF2 and Ulp1, and dKDM2 was found to be required for the ubiquitination of H2A by dRING-Psc (Lagarou *et al.* 2008). However, no obvious change in the overall levels of H2Aub in the *dKdm2* null mutant larvae was observed, which is consistent with a recent report showing that the dRING-L(3)73Ah complex plays the major role for the production of the bulk of ubiquitinated H2A: depletion of the PCGF protein L(3)73Ah, but not Psc and Su(z)2, in S2 cells significantly reduced the H2Aub level (Kahn *et al.* 2016).

The mammalian KDM2A (also known as FBXL11) and KDM2B (i.e., FBXL10) have a PHD domain; the PHD domain of the mammalian KDM2B can specifically bind to H3K4me3 and H3K36me2 (Janzer *et al.* 2012). It is interesting to note that the PHD domain is closely related to RING domains found in E3 ligases (Chasapis and Spyroulias 2009; Matthews *et al.* 2009). The PHD domain of the mammalian KDM2B has the E3 ubiquitin ligase activity, yet KDM2B alone could not ligate H2A with ubiquitin in the *in vitro* ubiquitination assay, suggesting the requirement of other proteins interacting with KDM2B for this activity (Janzer *et al.* 2012). It is interesting to note that the PHD domain is absent in KDM2B proteins in chicken, frog, zebra fish, and the invertebrate KDM2 proteins (Zheng *et al.* 2014). Taken together, these observations suggest that the major function of dKDM2 is to demethylate H3K36me2, and more work is needed to determine its role in targeting other epigenetic marks, particularly on specific promoters.

### dKDM2 regulates circadian rhythms

The circadian clocks enable animals to anticipate daily environmental changes and control most bodily functions including behavior and metabolism. In human osteosarcoma cells, it has been reported that depletion of human KDM2A shortens the circadian period, while its overexpression lengthens the period (Reischl and Kramer 2015). However, there is little evidence about the roles of KDM2 in circadian regulation at the organism level. Here we identified that in *Drosophila*, dKDM2 regulates circadian rhythmicity and period. Importantly, our *in vivo* data is consistent with previous studies in human cells: *dKdm2* nulls have a shortened period, while overexpression of dKDM2 lengthened circadian period (Fig 4, Table 1). In the preparation of this manuscript, we noticed that a recent study also observed that *dKdm2* mutants have shortened period (Shalaby *et al.* 2018). These results suggest that the roles of KDM2 in circadian rhythms are conserved among species.

How does dKDM2 regulate circadian period? To test whether dKDM2 controls the molecular pacemaker or circadian locomotor output, we have analyzed the levels of two critical pacemaker proteins in constant darkness CLK and PER. However, no obvious changes of these pacemaker proteins were observed, at least at the peak level. Although we cannot exclude potential effects on other circadian proteins, it is unlikely that dKDM2 affects the molecular pacemaker. Considering the expression of *36Y-GAL4* and the endogenous dKDM2 protein in peptidergic neurons including in the pars intercerebralis, it is possible that dKDM2 regulates circadian locomotor output pathway. Taken together, these results suggest that dKDM2 may regulate circadian clock through an unknown circadian output, instead of regulating the levels of the molecular pacemakers such as CLK and PER.

In conclusion, our results show that the major role of dKDM2 is the epigenetic regulation of circadian behavior, instead of normal development. Because small chemical inhibitors have been actively developed to specifically target KDMs in recent years (Hoffmann *et al.* 2012; Thinnes *et al.* 2014; Morera *et al.* 2016; Song *et al.* 2016; Tsai and So 2017), our results suggest that it will be important to determine whether the KDM2 inhibitors may affect the circadian clocks in both *Drosophila* and mammals. It will be necessary to perform detailed analyses of changes of epigenetic marks in neurons from *dKdm2* mutants, particularly on the genes involved in regulating circadian clocks and other behaviors in the future.

## Acknowledgement

We thank Paul Hardin and Weihua Li for helpful discussions, the Bloomington *Drosophila* Stock Center (NIH P40OD018537) for the fly strains, and the *Drosophila* Genomics Resource Center (NIH 2P40OD010949) for the pTWG vector. We thank Michael Rosbash, Paul Hardin for anti-PER and anti-CLK antibodies. The PDF antibodies, developed by Justin Blau was obtained from the Developmental Studies Hybridoma Bank, created by the NICHD of the NIH and maintained at the University of Iowa, Department of Biology, Iowa City, IA 52242. This work was supported by a grant from the NIH NIDDK (1R01DK095013 to JYJ), and YZ’s work was supported by the NIH COBRE Grant P20 (GM103650).

## Figure legends

**Fig. S1.**
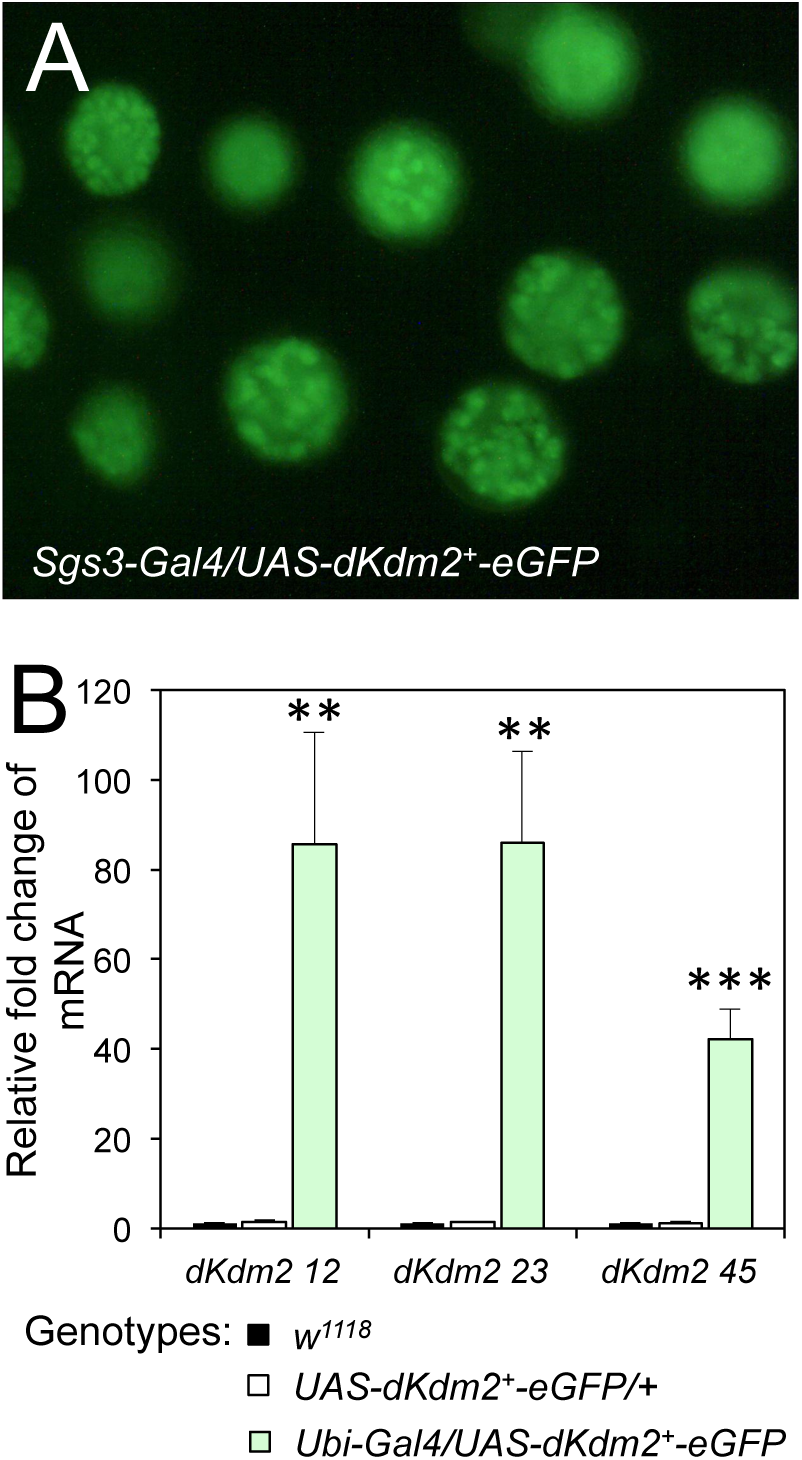
Characterization of the *UAS-dKdm2*^*+*^*-eGFP* line. (A) Nuclear localization of dKDM2-eGFP proteins driven by salivary-specific *Sgs3-Gal4* line. (B) Analysis of the mRNA levels of *dKdm2* in “*ubi-Gal4/UAS-dKdm2*^*+*^*-eGFP*” larvae using qRT-PCR. The regions uncovered by the primers are shown in Fig. 1A. The Student’s *t*-test was performed between the “*Ubi-Gal4/UAS-dKdm2*^*+*^*-eGFP*” line and *w*^*1118*^ (or the “*UAS-dKdm2*^*+*^*-eGFP /+*” line), and the * above green bars represents both two set of the *t*-tests.

**Fig. S2.**
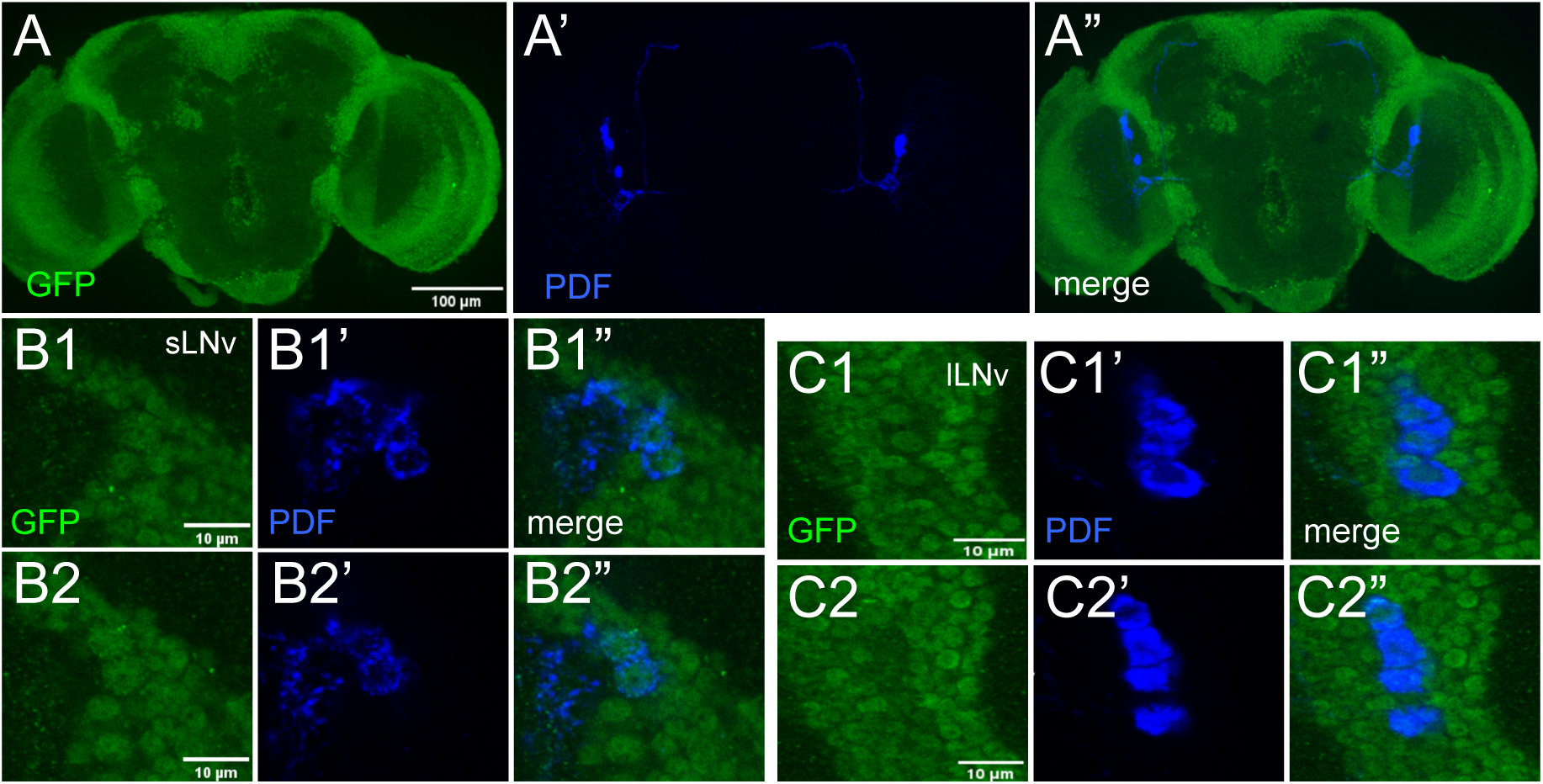
Expression of the endogenous dKDM2 protein tagged with eGFP in adult brain. (A) Representative confocal image of dKDM2-eGFP (green), stained together with antibody against PDF (A’, blue), and the merged image is shown in A”. Scale bar in (A): 100 µm. (B) Expression of dKDM2 in sLNv, B1 and B2 are two successive focal planes. (C) Expression of dKDM2 in lLNv, C1 and C2 are two successive focal planes. Scale bars in B1/B2 and C1/C2: 10 µm.

**Fig. S3.**
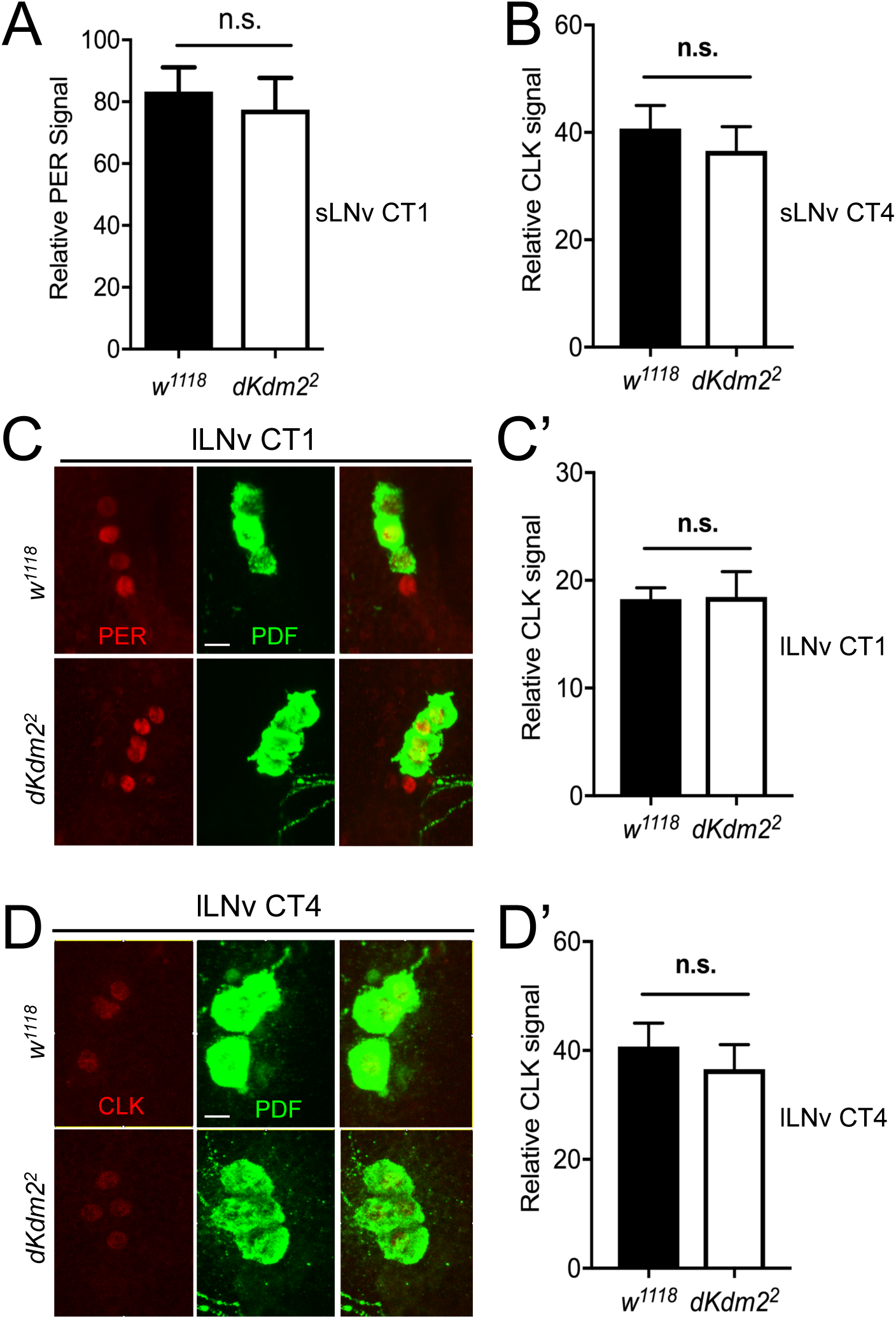
Quantification of PER and CLK levels in *dKdm2* mutants. (A, B) Bar graph shows the abundance of PER in sLNv CT1 and CLK in sLNv CT4. (C) Expression of PER (in red) in lLNv CT1, co-stained with anti-PDF (in green), and the results are quantified and shown in C’. (D) Expression of CLK (in red) in lLNv CT4, co-stained with anti-PDF (in green), and the results are quantified and shown in D’.

